# qDSB-Seq: quantitative DNA double-strand break sequencing

**DOI:** 10.1101/171405

**Authors:** Yingjie Zhu, Anna Biernacka, Benjamin Pardo, Norbert Dojer, Romain Forey, Magdalena Skrzypczak, Bernard Fongang, Jules Nde, Raziyeh Yousefi, Philippe Pasero, Krzysztof Ginalski, Maga Rowicka

## Abstract

Sequencing-based methods for mapping DNA double-strand breaks (DSBs) allow measurement only of relative frequencies of DSBs between loci, which limits our understanding of the physiological relevance of detected DSBs. We propose quantitative DSB sequencing (qDSB-Seq), a method providing both DSB frequencies per cell and their precise genomic coordinates. We induced spike-in DSBs by a site-specific endonuclease and used them to quantify labeled DSBs (e.g. using i-BLESS). Utilizing qDSB-Seq, we determined numbers of DSBs induced by a radiomimetic drug and various forms of replication stress, and revealed several orders of magnitude differences in DSB frequencies. We also measured for the first time Top1-dependent absolute DSB frequencies at replication fork barriers. qDSB-Seq is compatible with various DSB labeling methods in different organisms and allows accurate comparisons of absolute DSB frequencies across samples.

## Introduction

There is tremendous interest in precisely measuring DNA double-strand breaks (DSBs) genome-wide since such measurement can give key insights into DNA damage and repair, cancer development^1^, radiation biology, and also increasingly popular genome editing techniques^2^. Starting with our BLESS method^3^, several high-resolution and direct methods to label DSBs genome-wide have recently been developed^4-7^, which have opened up new possibilities for sensitive and specific detection of DSBs. For example, BLESS was applied in identifying the on-target and off-target cutting sites of Cas9 endonuclease^8^ and studying DSB repair^9^. However, we still lack an effective strategy to both precisely detect DSB distribution genome-wide and quantify their absolute frequencies per cell, which is crucial to assess physiological relevance of detected DSBs. Immunofluorescence microscopy in combination with γ-H2AX and 53BP1 antibodies was used to count breaks per cell^10^, but does not allow determining their precise locations. Moreover, counting discrete nuclear foci is an imprecise way to estimate DSB numbers per cell both due to DSB clustering and limited specificity of antibodies. Quantitative Polymerase Chain Reaction (qPCR) based methods can estimate absolute break frequency but only at selected loci^11^. An approach was developed recently to quantify breaks globally based on amount of radiolabeled DNA and locally based on DNA break immunocapture^12^, but its accuracy in detecting physiological DSBs was not tested. BLISS^7^ quantifies DSBs by utilizing unique molecular identifiers (UMIs) to identify unique DSB ends and counting cells in the sample. BLISS is designed for detecting DSBs in samples with low number of cells and thus shares challenges of single-cell sequencing, such as low genome coverage and over-amplification. Moreover, employment of UMIs is challenging. Short UMIs may lead to UMI collisions^13^ (i.e. observing two reads with the same sequence and the same UMI barcode but originating from two different genomic molecules), especially in case of DSBs enriched in specific locations. Long UMIs may interfere with primer sequence binding and accumulate sequencing errors, which may lead to severe overestimation of DSBs^14^.

This lack of a general method and computational solution to simultaneously determine DSB frequencies per cell and their precise genomic loci limits our understanding of the physiological relevance of observed DSBs and hinders comparisons between experiments. Here, we propose quantitative DSB sequencing (qDSB-Seq), an approach that allows measuring DSB frequencies per cell genome-wide, and a computational solution to achieve accurate quantification. Our approach relies on inducing spike-in DSBs by a site-specific endonuclease, which are used to quantify DSBs detected by a DSB labeling method e.g. i-BLESS^15^ and can be combined with any DSB labeling technique. We present a comprehensive validation and several applications of qDSB-Seq: quantifying DSBs induced by a radiomimetic drug, occurring during replication stress and caused by natural replication fork barriers.

## Results

### qDSB-Seq implementation, computational method and validation

qDSB-Seq is a combination of genome-wide high-resolution DSB-labeling (i-BLESS^15^, BLESS^3^, END-seq^6^, etc.) and inducing DSBs (spike-ins) in pre-determined loci using a site-specific endonuclease (**Fig. 1a-c**). Quantification is based on an assumption (verified below) that the number of labeled reads at a given genomic locus resulting from DSB sequencing is proportional to the underlying DSB frequency (proportionality coefficient *α* in **Fig. 2a**).

**Figure 1.**
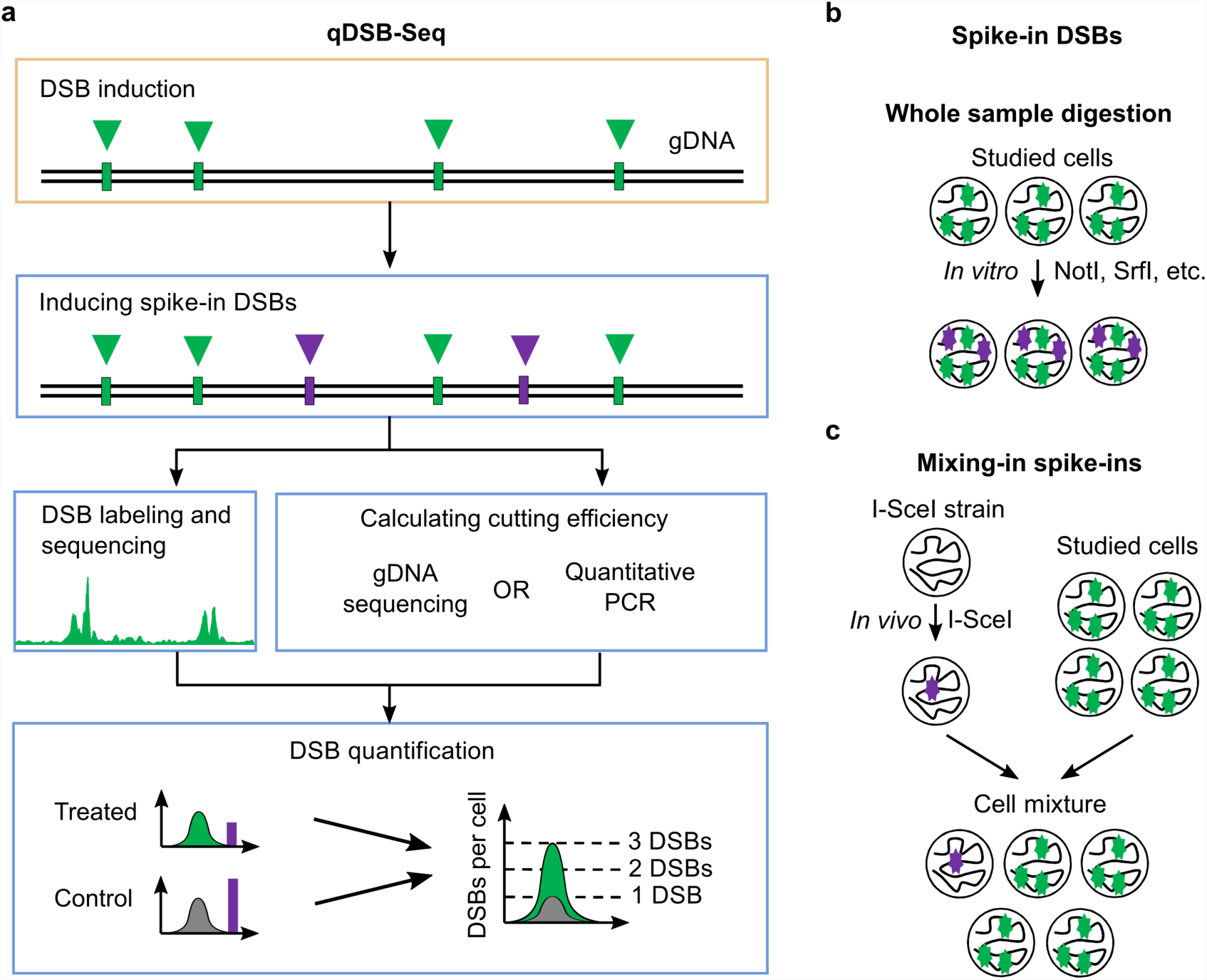
qDSB-Seq method. **(a)** In qDSB-Seq protocol after DSBs induction, cells are treated with a restriction enzyme to introduce site-specific, infrequent DSBs (spike-ins). Next, DSBs are labeled (using e.g. i-BLESS) and sequenced. Simultaneously, gDNA sequencing (or qPCR) is performed and used to estimate the cutting efficiency of the enzyme, and thus frequency of induced DSB spike-ins, which is then used to quantify the absolute DSB frequency (per cell) of studied DSBs in the sample (**Methods**). **(b-c)** Spike-in DSBs were induced in two different ways: (**b**) the studied cells were digested using the NotI, SrfI, AsiSI, or BamHI restriction enzyme *in vitro*; (**c**) cells expressing the restriction enzyme I-SceI *in vivo* (the I-SceI strain) were mixed with the studied cells.

**Figure 2.**
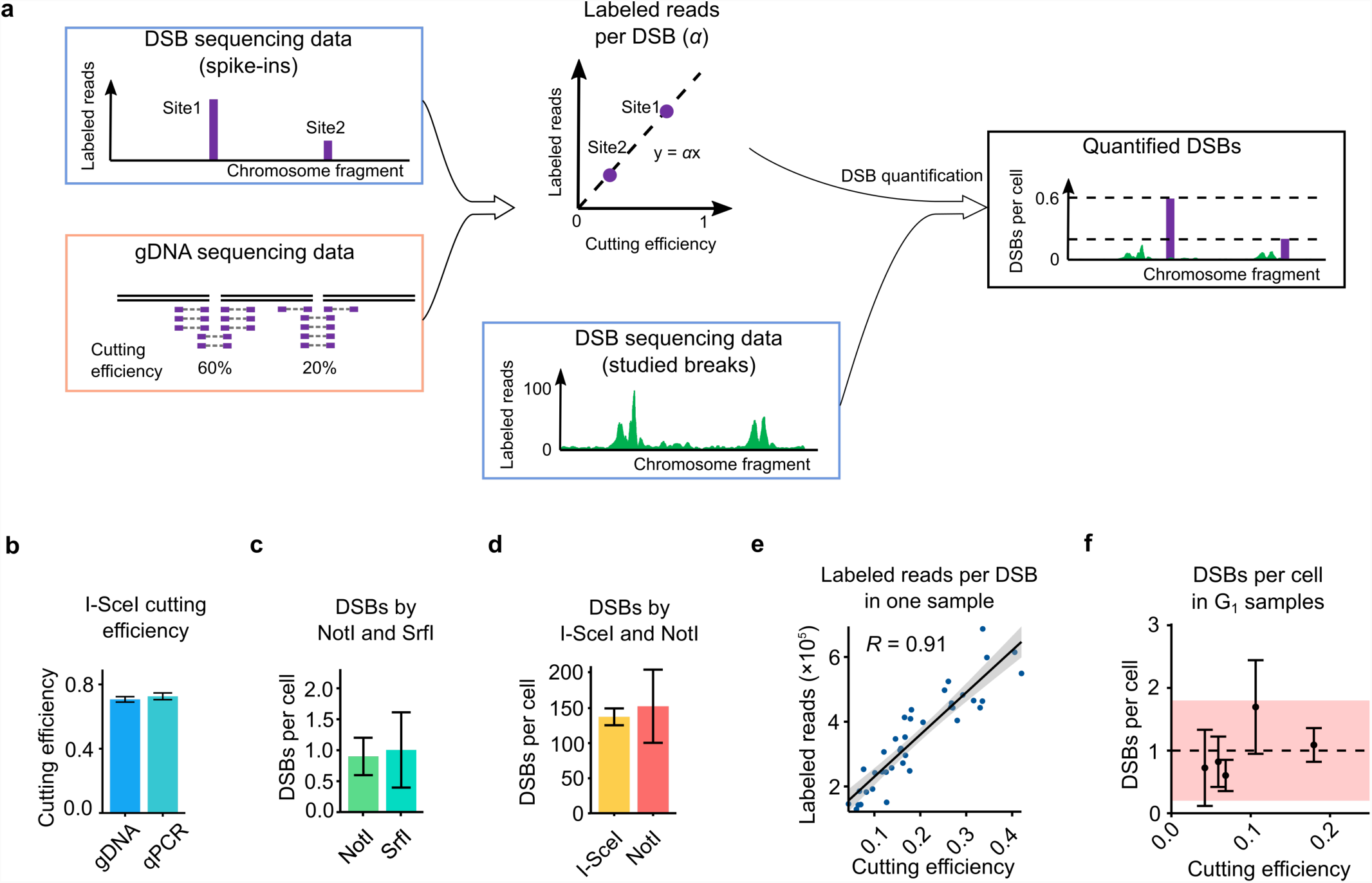
qDSB-Seq computation and validation. **(a)** Computation of the labeled reads per DSB and DSBs per cell. The ratio of the labeled reads and cutting efficiency at enzyme cutting sites was calculated and then used for DSB quantification in the studied genomic loci. **(b)** The estimation of I-SceI cutting efficiency based on gDNA sequencing and qPCR (error was calculated from technical replicates). **(c-d)** Dependence of qDSB-Seq quantification on the restriction enzyme used. DSB levels obtained for **(c)** untreated WT G_1_ phase cells, and for **(d)** HU-treated WT S phase cells quantified using NotI and SrfI digestion *in vitro* and I-SceI digestion *in vivo* (errors of the estimated DSB frequencies were calculated as described in **Methods**). **(e)** Correlation between the number of labeled reads at cutting sites and their cutting efficiencies in an untreated G_1_ phase sample, digested with NotI enzyme with average cutting efficiency of 18%. *R*: Pearson correlation coefficient. **(f)** Quantification of DSBs in untreated G_1_ phase cells. The dashed lines and the stripes are the mean value and 95% confidence interval, respectively. Mean ± s.d. is shown for each sample.

To estimate this coefficient *α*, we induce spike-in DSBs at pre-determined genomic loci and, relying on knowledge of their exact genomic locations, quantify their frequency using genomic DNA sequencing (gDNA) or qPCR (**Fig. 1a, Fig. 2a**). The spike-in DSBs are created by digestion with a restriction endonuclease before DSB labeling (**Fig. 1b,c**). Next, the frequency of induced spike-in DSBs, *B*_*cut*_, is calculated from enzyme cutting efficiency, *f*_*cut*_, that is calculated from gDNA sequencing data based on numbers of cut and uncut DNA fragments covering cutting sites in gDNA (**Fig. 2a, Methods**), or qPCR data (**Supplementary Fig. 1, Methods)**.

Finally, the absolute frequency of studied DSBs, *B*_*studied*_, is estimated from DSB sequencing data:

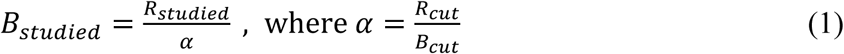

and *R*_*studied*_ and *R*_*cut*_ are the numbers of labeled reads originating from studied DSBs and from enzyme cutting sites (spike-ins), respectively, and *B*_*cut*_ ∼ *f*_*cut*_.

### Reproducibility and accuracy of cutting efficiency estimation

The number of labeled reads per DSB (coefficient *α*) which is used for the final DSB quantification, as explained above, is computed from enzyme cutting efficiency, *f*_*cut*_ (**Equation (1), Methods**). Therefore, to calculate *α* accurately, we need to be able to estimate enzyme cutting efficiency accurately. Commonly qPCR is used for precise measurement of cutting efficiencies, however, this technique is inconvenient to use for multiple cutting sites. Thus, we propose to use gDNA sequencing to determine spike-in cutting efficiencies **(Fig. 2a, Methods)**. To verify the accuracy and reproducibility of our proposed approach, we treated immobilized and deproteinized yeast DNA with NotI enzyme and compared cutting efficiencies at its recognition sites calculated using gDNA sequencing data and qPCR. The cutting efficiencies for the selected NotI cutting site were highly consistent: 61% for gDNA sequencing and 62% for qPCR. To examine if our approach can also be applied to breaks introduced *in vivo*, which can be subjected to repair and resection, we used a yeast strain engineered to produce a single site-specific DSB by I-SceI endonuclease *in vivo.* Cutting efficiencies calculated based on gDNA sequencing and based on qPCR (**Supplementary Fig. 1, Methods**) were again very consistent: 71% and 73%, respectively (**Fig. 2b)**. We therefore conclude that our method of estimating enzymes cutting efficiency based on gDNA sequencing yields accurate and precise results.

### Dependence of quantification on enzyme choice and types of breaks induced

DSBs occurring *in vivo* are subject to DNA damage repair and therefore might be labeled with different efficiencies than breaks induced *in vitro*. Moreover, different types of double-stranded DNA ends (blunt or sticky) could also be detected more or less efficiently by a given DSB labeling method. We therefore asked whether any restriction enzyme and any manner of digestion can be applied to create spike-in DSBs that would lead to accurate quantification. First, to test if restriction enzyme choice or the types of double-stranded DNA ends influences our quantification results, we determined the spontaneous DSB frequencies in yeast G_1_ phase cells using NotI or SrfI spike-ins, which create sticky and blunt ends, respectively. The number of spontaneous breaks in G_1_ phase cells estimated using these enzymes was consistent: 0.9 ± 0.3 DSBs per cell for NotI spike-in and 1.0 ± 0.6 DSBs per cell for SrfI spike-in (**Fig. 2c**). Then, to test if the results are affected by the manner of digestion, we compared DSB estimations based on quantification using NotI (5’ overhangs) *in vitro* digestion and I-SceI (3’ overhangs) *in vivo* digestion in HU-treated wild-type cells (described below). Again, results were highly similar: 137 ± 12 and 153 ± 52 DSBs per cell (**Fig. 2d**). In conclusion, qDSB-Seq provided consistent results in all tested cases irrespective of the restriction enzyme used, types of DNA ends created by that enzyme, or the manner of digestion.

### Dependence of accurate quantification on adequate cutting efficiency

For accurate quantification of studied DSBs, it is necessary that the relationship between the number of labeled reads and DSB frequencies at different genomic locations is linear (**Equation (1), Fig. 2a**). This relationship could be affected by the frequencies of spike-in DSBs, which is determined by an enzyme cutting efficiency. Therefore, we asked whether any frequency of induced spike-in DSBs (i.e. any enzyme cutting efficiency) can be employed. To test the influence of enzyme cutting efficiency on the quantification results, we performed 35 digestions for 25 samples using enzymes with multiple cutting sites (NotI, SrfI, AsiSI, and BamHI) and then tested the linear relationship between the labeled reads and cutting efficiencies for each digestion using Pearson Correlation Coefficient. We observed that strong correlation (*R* > 0.5) (e.g. **Fig. 2e**) was always achieved for cutting efficiencies between 12% and 62% (**Supplementary Fig. 2**, **Supplementary Table 2**) and for some lower cutting efficiencies (4-12%). However, for the extreme cutting efficiencies (higher than 84% or lower than 4%) the correlation was always weak (**Supplementary Fig. 3**). In such cases, the number of observed cut or uncut fragments was low, making our estimates less accurate, which likely decreased the correlation. Moreover, small variations in *f*_*cut*_ between sites contributed to the decreased correlation (**Supplementary Fig. 3**). Additionally, in samples for which digestion efficiencies are very high, the elevated level of reads at spike-in sites (> 75%) (**Supplementary Table 1**) can potentially disrupt (due to low initial sequence diversity) Illumina sequencing^16^. Taken together, we conclude that adequate cutting frequencies (4% to 84%) lead to a constant ratio between the labeled reads and the cutting efficiencies for accurate quantification.

### Stability of estimation of DSB frequencies per cell

We next asked whether our method generates reproducible results. To test this, we calculated DSB frequencies in untreated G_1_ cells based on different spike-ins. In spite of the various enzymes used (NotI, SrfI) we obtained a very consistent number of DSBs (**Fig. 2f, Supplementary Fig. 4**, **Supplementary Table 1**). Based on our calculations the frequency of spontaneous DSBs in untreated G_1_ wild-type cells is 1.0 ± 0.4 DSBs per cell, both the average and the range (0.6-1.7 DSBs per cell) are consistent with previous studies^17, 18^ (**Supplementary Table 1**). Further, we quantified DSBs based on the individual cutting sites in each of these samples. The variation of the DSB quantification results depending on the individual cutting sites used was lower than the average value (**Supplementary Table 1**). Similarly, in *pif1* mutants, where stability of some DNA secondary structures is affected and we observed increase in DSB numbers related to G-quadruplex^15^ structures, we obtained average DSB number 2.1 DSBs per cell. DSB quantification was consistent between the samples (s.d. 0.3 DSBs per cell) (**Supplementary Fig. 4**, **Supplementary Table 1**).

## Applications of qDSB-Seq

### Quantification of DSBs induced by a radiomimetic drug, Zeocin

Some DSB-inducing agents affect only particular sequences and structures, while others cause DNA damage throughout the genome, e.g. irradiation. As DSB sequencing data inform only about read distribution in the genome and is primarily used to identify regions enriched in reads, even very large but global DSB induction will be undetectable using typical normalization methods, e.g. normalization to the background. Therefore, to test application of qDSB-Seq to such a challenging case, we used the radiomimetic agent Zeocin^19^, a member of the bleomycin drug family. After performing DSB sequencing, no apparent difference in raw read counts between Zeocin-treated (ZEO) and untreated G_1_ phase (G_1_) cells was observed (**Fig. 3a, Supplementary Fig. 5**). In contrast, after quantification (using qDSB-Seq with NotI spike-in) we concluded that 1.1 ± 0.3 DSBs/cell were present in the G_1_ sample and 7.4 ± 1.7 in ZEO, indicating that Zeocin induced 6.3 ± 2.0 DSBs per cell. Strikingly, Zeocin significantly increased the number of DSBs (1.7- to 13-fold) in 99.8% of 5 kb genomic intervals (p-value < 2e-12, hypergeometric test, **Methods**).

**Figure 3.**
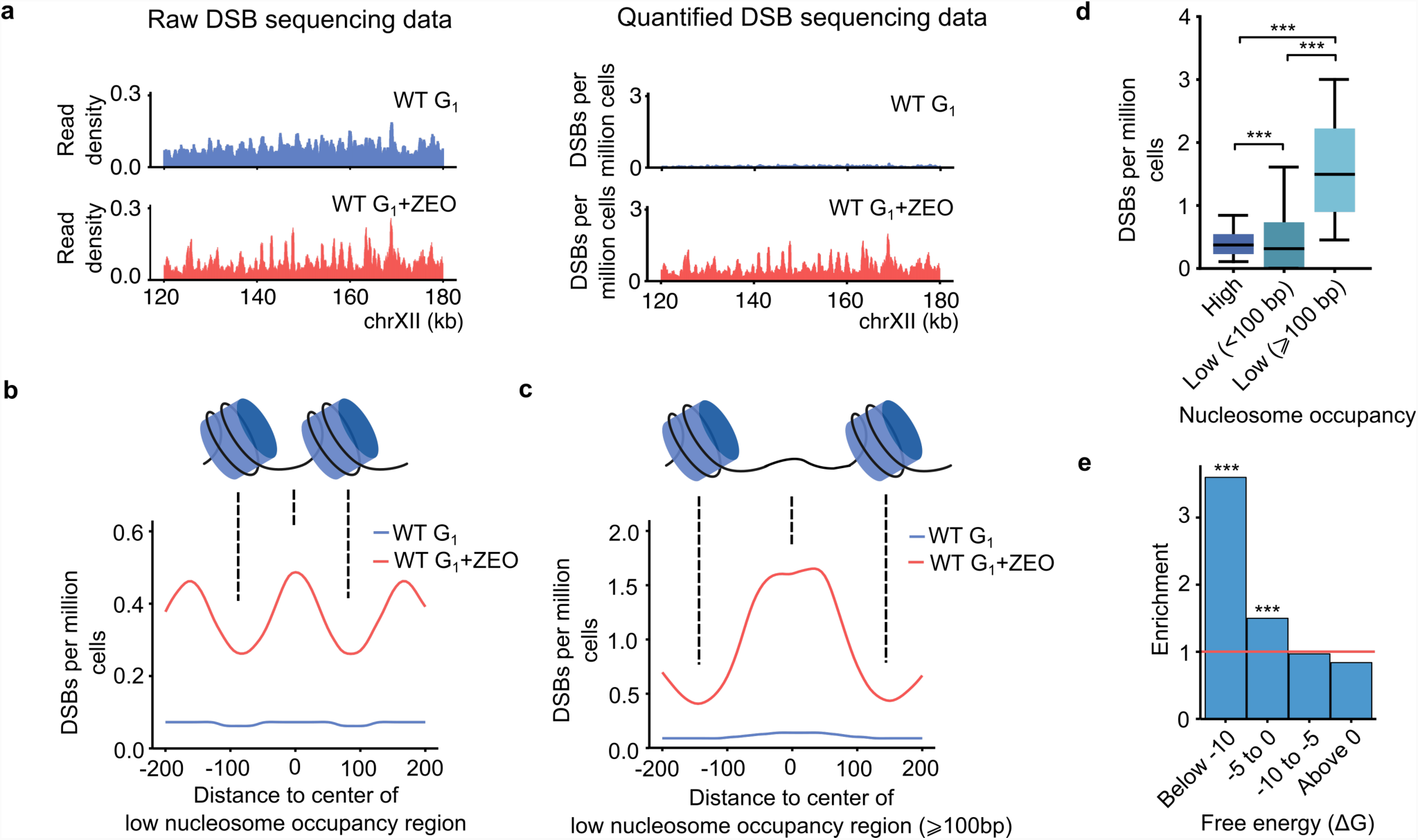
Quantification of Zeocin-induced DSBs. **(a)** Raw read density and quantified DSB density (500 bp sliding window with 50 bp step) in a representative fragment of chromosome XII for untreated and Zeocin-treated wild-type G_1_ phase cells. Raw read density was normalized to the total number of reads. **(b-c)** Density of Zeocin-induced DSBs in **(b)** all and **(c)** ≥100 bp low nucleosome occupancy regions. Nucleosome locations from Lee *et al.*^20^ were used, DSB densities, expressed as DSBs per million cells, were calculated in a 50 bp sliding window with a 5 bp step. **(d)** Comparison of DSB densities in high nucleosome occupancy regions (High) and low nucleosome occupancy regions (Low). **(e)** Enrichment of Zeocin-induced DSBs in regions prone to form very stable DNA secondary structures (e.g. hairpins), as defined by free energy in a 50 bp sliding window as described in **Methods**. Zeocin-induced DSBs were defined as regions with significant enrichment of DSB-labeled reads in ZEO sample compared with G_1_ phase control, as identified using Hygestat_BLESS. Enrichment analysis was performed using hygestat_annotations (**Methods**). *P* values: * *P* < 0.05, ** *P* < 0.01, *** *P* < 0.001.

Interestingly, we observed that Zeocin-induced DSBs are especially enriched (3.0-fold) in nucleosome-depleted regions (NDR) and reduced (0.4-fold) in nucleosome-protected regions (both *p* < 10^-3^, permutation test, **Methods**). Specifically, DSBs in the Zeocin-treated sample occur 1.8 times as often between predicted nucleosome positions^20^ as within nucleosomes (**Fig. 3b**). Moreover, the preference for DSB location between nucleosomes is even higher (4.1-fold) for long (> 100 nt) NDR regions (**Fig. 3c,d**). However, we do not observe a 10 bp periodicity corresponding to the rotational positioning of the DNA helix on the nucleosome. These results are consistent with previous findings that Zeocin-induced cleavage is most suppressed in nucleosome-bound DNA and that this suppression is not dependent on inaccessibility of the minor groove, but is caused by inability of the nucleosome-bound DNA to undergo a conformational change that is required for Zeocin binding^21^. Zeocin-induced DSBs are also enriched in DNA regions capable of forming very stable DNA secondary structures (**Fig. 3e**), including G-quadruplexes (G4s)^22^. Further studies will be necessary to elucidate this phenomenon. Nevertheless, increased DNA damage on G4 structures could be related to nucleosome remodeling on G4s^23^, consistent with our finding that Zeocin prefers to cleave nucleosome-free DNA.

### Quantification of DSBs induced by replication stress

We next used qDSB-Seq to quantify replication-associated DSBs under hydroxyurea-induced replication stress (**Fig. 4a**). Hydroxyurea (HU) inhibits ribonucleotide reductase, resulting in decreased dNTP levels and subsequent replication fork stalling and the slowing down of S phase^24^. Without the protection of replication checkpoints, stalled forks may undergo catastrophic collapse at high concentration or prolonged HU treatment^25^, such as we used.

**Figure 4.**
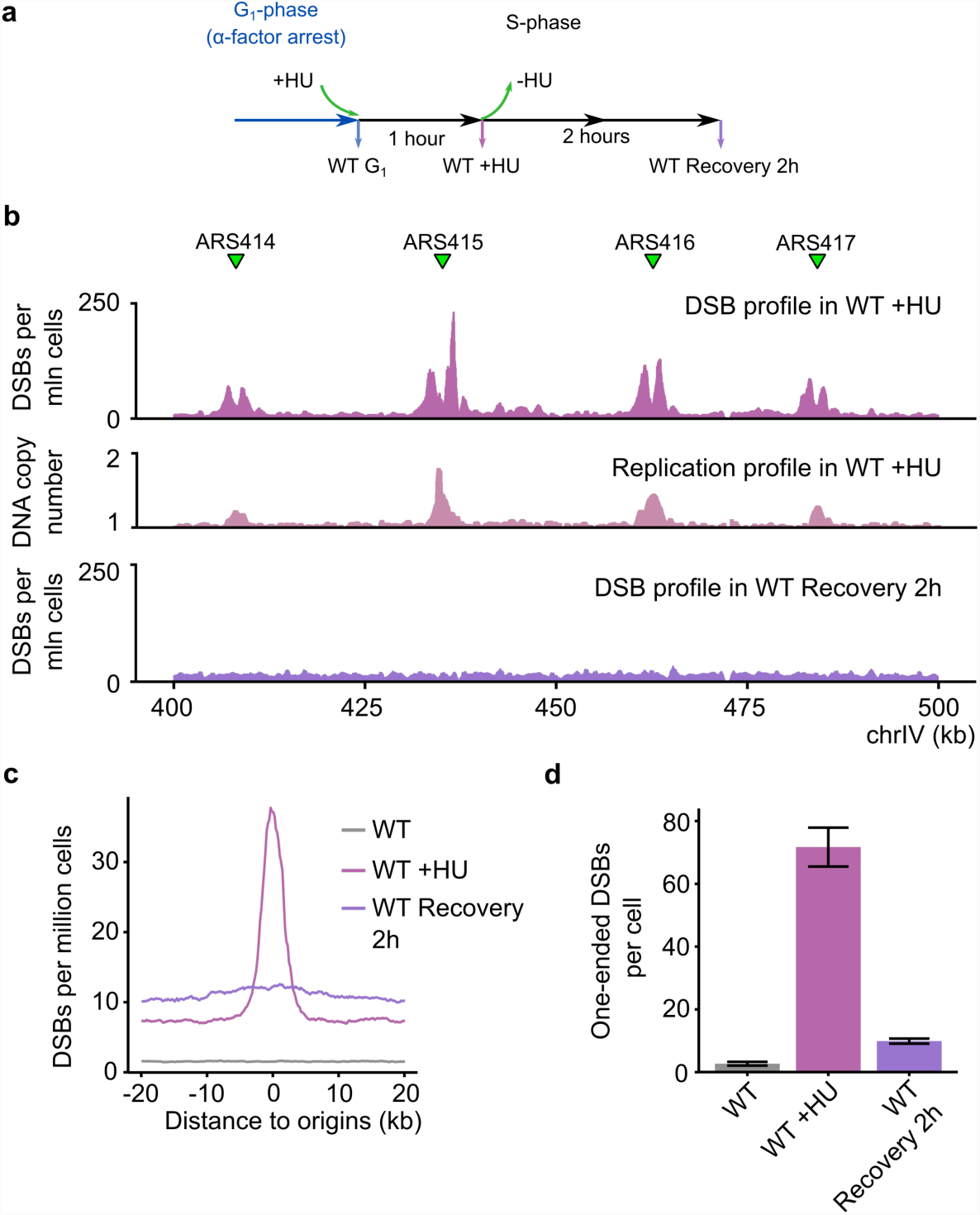
Quantification of replication-associated DSBs. **(a)** Schematic representation of HU experiments. Cells were arrested in G_1_ phase with *α*-factor, treated with HU before release to S phase, harvested after 1 hour or resuspended in fresh medium and harvested 2 hours after removal of HU. **(b)** Example of quantified DSB data from HU-treated wild-type and 2-hour recovery cells. Replication origins are marked with green triangles, absolute frequencies of DSBs for a fragment of chromosome IV are shown in a million cells. As a control, replication profile (values of DNA copy number) in WT +HU sample is shown, for which the number of gDNA reads in a 500 bp window in WT +HU sample was normalized by G_1_ sample. **(c)** Meta-profile of DSBs around active replication origins under HU treatment, defined as 144 origins with firing time < 25 min (early origins, firing time according to Yabuki *et al.*^39^}). Median of DSB densities, expressed as DSBs per million cells in 2 kb window around each early origin, was calculated, the background was removed as described in **Methods**. **(d)** Quantification of one-ended DSBs. Errors of the estimated one-ended DSB frequencies were calculated as described in **Methods**.

Using NotI spike-in, we observed that one hour treatment with 200 mM HU induced on average 137.6 ± 12.0 DSBs per cell in wild-type yeast cells (WT +HU sample), which represents a 9-fold increase relative to untreated S phase cells (15.4 ± 3.2 DSBs per cell). The detected breaks showed a clear replication-related pattern: a significant enrichment of DSB signal around replication origins (**Fig. 4b,c**). To further analyze the HU-induced DSBs we classified them into two-ended DSBs and one-ended DSBs (**Supplementary Fig. 6)**. Two-ended DSBs arise when two strands of DNA double helix are damaged (by i.e. endonucleases, radiation or chemical compounds), while broken replication forks result in one-ended DSBs. We identified one-ended DSBs using our method based on comparing the number of reads between Watson and Crick strands (**Supplementary Fig. 6**, **Methods**) and discovered that among all DSBs detected in HU-treated WT cells 71.7 ± 6.2 DSBs were one-ended (**Fig. 4d**). Of those, 85% (60.6 ± 5.2 DSBs) were located within +/-10 kb regions of active origins, resulting in an average of 0.4 one-ended DSB (broken fork) per origin (**Fig. 4d**). Such one-ended DSBs would not be detected by some other DSB detecting methods, such as pulse-field gel electrophoresis, which explains some earlier reports that wild-type yeast cells are not sensitive to HU^25^. The observed one-ended DSBs might correspond to broken forks resulting from transient DNA breaks occurring on the leading strand, as reported by Sasaki *et al*^26^. In agreement with this theory, we discovered that two hours after removal of HU, the number of one-ended DSBs decreased dramatically (by 86%) (**Fig. 4d**), indicating that replication-associated DNA damage present during HU treatment is not permanent.

### Quantification of DSBs at ribosomal replication fork barriers

Replication fork barriers (RFBs) are natural barrier that blocks replication forks to protect nearby, highly expressed rRNA genes from collisions between transcription and replication complexes^26, 27^ (**Fig. 5a**). DSBs occurring at the ribosomal replication fork barriers (RFBs) have been observed using Southern blot in the budding yeast^28-31^. However, precise frequencies and genomic locations of these DSBs were not established due to lack of a quantitative and sensitive DSB detection method^26^. Using qDSB-Seq, here we both precisely quantified DSB frequencies near RFBs and identified their genomic coordinates.

**Figure 5.**
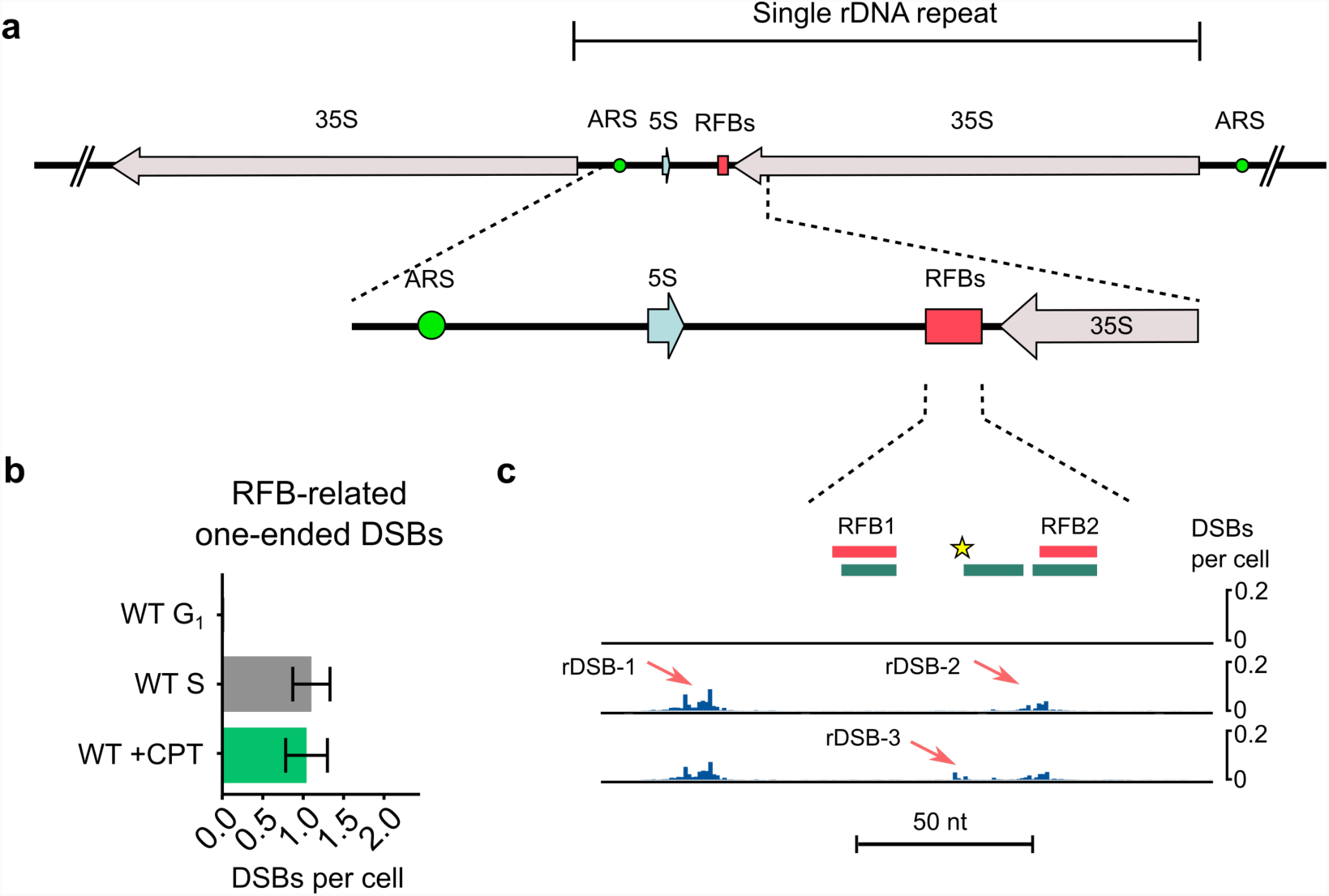
DNA double-strand breaks at replication fork barriers. **(a)** Scheme of Replication Fork Barriers (RFBs) at yeast rDNA locus. **(b)** The total number of RFB-related one-ended DSBs (peaks as defined in panel **c)** calculated from the difference of Watson and Crick strand reads (**Methods**); **(c)** Quantified DSBs signal in RFB region. RFB1 and RFB2 are indicated by the red boxes on the top. The green boxes mark Fob1 protein binding sites mapped *in vitro*. The yellow star indicates Top1 cleavage site. The red arrows point out the observed ribosomal DSB sites, rDSB-1, rDSB-2, and rDSB-3.

It was reported that Fob1 proteins bound to an RFB site block replication fork progression, resulting in generation of a one-ended DSBs^30^. Indeed, in unperturbed S-phase cells, we observed 1.1 DSBs per cell (0.0055 DSBs per rDNA repeat) on rDSB-1 and rDSB-2 sites upstream of RFB1 and RFB2 (two closely spaced RFB loci) (**Fig. 5b,c and Supplementary Table 3**). As expected, we did not detect any DSBs at these sites in G_1_-arrested cells confirming that the observed DSBs at RFBs are replication-dependent.

It was previously shown that Top1 in the presence of Fob1 specifically cleaves defined sequences in the RFB region^32^. When we inhibited the religation step of Top1 by adding 100 μM camptothecin (CPT) for 45 min treatment, we observed a CPT-dependent DSB site (rDSB-3), exactly at the same location as the previously identified Top1-dependent cleavage site (**Fig. 5c**). In addition, this site also colocalizes with a Fob1 binding region, in agreement with a previous discovery that the recruitment and stabilization of Top1 requires the binding of Fob1 protein^32^. Our quantification shows the DSB frequency at rDSB-3 site was 0.1 DSB per cell, lower than at rDSB-1 and rDSB-2. Finally, our results agree with previous work^26^ in which approximately one DSB arises in an rDNA array during replication in a yeast cell (**Fig. 5b**); such low frequencies are caused by recombination in the rDNA array^26^. Based on the results above, qDSB-Seq fills the need to enable detection of these rare breaks at replication fork barriers and allowed us for the first time to quantify the frequency of cleavage of Topoisomerase 1 (Top1) at RFBs.

## Discussion

We propose qDSB-Seq, a general framework that allows estimating both absolute DSB frequencies (per cell) and their precise genomic coordinates. qDSB-Seq combines a DSB-labeling method with a quantification technique; quantification is achieved by inducing easy-to-measure spike-in DSBs via restriction enzyme digestion.

Due to increasing evidence of a relationship between emergence of DSBs and human diseases such as cancer^1^, there is growing interest in precise detection of DSBs. Several general genome-wide methods for detection of DSBs with single-nucleotide resolution have recently been developed^3-6^, however their usefulness is limited because they only allow comparison of DSB levels between genomic loci within the same sample.

Normalization to the total number of reads is often employed to enable comparison between different samples, but this method is not always applicable. For example, it cannot be used if DSBs are induced throughout the whole genome or if the DSB background varies, which is common^33^. Therefore, in case of agents that create such DSB patterns, e.g. by irradiation or radiomimetic drugs, data normalized to the total reads number will not reveal global induction of breaks as shown in **Fig. 3a**. In contrast, our approach allows not only estimation of relative increases of DSB signal between samples (regardless of signal distribution), but also quantification of absolute DSB numbers per cell. For example, we discovered that 1 hour treatment with 100 µg/ml Zeocin results in 6.7-times increase in DSBs, namely from 1.1 ± 0.3 to 7.4 ± 1.7 DSBs per cell. Additionally, we discovered that Zeocin significantly increases DSB levels in 99.8% of 5kb genomic intervals, but with differences in ratios: from 1.7- to 13-fold. qDSB-Seq opens up new possibilities in studying the impact of DSB inductors or gene mutations on genome instability, i.e. it may potentially allow determining the outcomes of different doses of anticancer drugs in healthy and tumor cells. Moreover, qDSB-Seq allows assessing DSB frequencies not only for the whole genome, but also for a specific locus. For instance, using our approach, for the first time we quantified changes of DSB frequency at RFBs between wild-type and CPT-treated cells, thus revealing the frequency of Top1-dependent DSBs in RFB region.

Key innovation of qDSB-Seq is spike-in DSBs used for normalization. Such spike-in DSBs can be introduced both *in vivo* and *in vitro*; each manner of digestion has its strengths and weaknesses. *In vivo* digestion requires organism-specific constructs, such as the I-SceI yeast strain we used, while *in vitro* digestion can be applied to any organism. Moreover, for *in vitro* digestion, since spike-in DSBs are never repaired and thus there are no resected DNA ends. Resected DNA ends may result in spike-in related reads located up to several kilobases from the cutting sites, which may complicate data analysis. On the other hand, for *in vivo* digestion it is possible to determine enzyme cutting frequency before addition of spike-in cells to the sample of interest, which facilitates obtaining final cutting efficiency in the desired range by selecting desired mixing proportions. *In vivo* digestion can be also used to study the DNA damage response in systems such as DivA^34^.

Enzyme cutting efficiency is a key parameter influencing qDSB-Seq accuracy. As shown above, using extremely low or high cutting efficiencies may result in inaccurate quantification results, while within an adequate range (4% to 84%), the number of labeled reads per DSB (proportionality coefficient *α*) remains constant, which allows for consistently accurate quantification. If spike-in DSBs are introduced *in vivo*, to achieve desired cutting efficiency one needs to mix in appropriate proportions cells in which full digestion (or digestion with known efficiency) was performed with the studied cells. In case of *in vitro* digestion, the studied cells should be treated with a dose of an enzyme much lower than recommended for full digestion. The enzyme cutting efficiency can be then estimated by performing qPCR and, if needed, the dose can be adjusted before sequencing.

To facilitate choice of a restriction enzyme for qDSB-Seq experiments we provide lists of restriction enzymes sorted according to their cutting efficiencies per Mb in the yeast, human, mouse and fruit fly genomes (**Supplementary Table 4**), as well as Genome-wide Restriction Enzyme Digestion STatistical Analysis Tool, GREDSTAT, at http://bioputer.mimuw.edu.pl:23456. Enzymes with multiple cutting sites should yield best quantification results, since estimation of the enzyme cutting frequency will be less influenced by a potential local bias. Constructs with a single enzyme cutting site, such as the I-SceI strain we employed, allow convenience of using qPCR to determine an enzyme cutting frequency. Therefore, for enzymes with multiple cutting sites, we developed a method to estimate enzyme cutting efficiency from gDNA sequencing data, and proved its accuracy by comparing with qPCR results. On the other hand, usage of rare cutting enzymes is preferable, since they allow for optimal cutting efficiencies at individual sites without unnecessarily increasing percentage of spike-ins in total reads. There is no benefit to using a higher spike-in percentage than necessary; high spike-in percentages, especially exceeding 30-50% of total reads, may cause quality issues with Illumina sequencing^16^. Unlike enzyme cutting efficiency, percentage of spike-in reads cannot be determined before sequencing, since it depends both on enzyme cutting efficiency and number of DSBs present in the data. Therefore, if there is a probability that high level of spike-ins may be achieved unintentionally (e.g. during pilot experiments), we recommend using our modified protocols for generation of high-quality sequencing data from low-diversity samples^16^.

qDSB-Seq is compatible with any DSB labeling technique, but will also share limitations of the used method. For example, we tested that the type of generated DNA ends will not determine quantification results when using i-BLESS for DSB labeling. However, as we discussed in^15^, some DSB sequencing technologies cannot detect all types of DNA ends. Therefore, qDSB-Seq, when used in combination with such technology, will also exhibit bias in quantifying DSBs with these types of DNA ends.

When interpreting qDSB-Seq results, it is important to keep in mind that qDSB-Seq relies on sequencing data derived from a population of cells. Therefore, it only yields an average number of DSBs per cell, which may or may not be representative of a typical single cell. This problem can be solved by combining qDSB-Seq with a complementary method, giving insight into population-distribution of DSBs, as we proposed elsewhere^33^.

In summary, qDSB-Seq is a novel approach, which allows absolute DSB quantification genome-wide and accurate cross-sample comparison and can be applied to any organism, for which a DSB labeling method is available. qDSB-Seq relies on a key innovation, using spike-in DSBs induced by a restriction enzyme for normalization. Using qDSB-Seq, we quantified the numbers of DSBs induced by a radiomimetic drug and replication stress; measured for the first time Top1-dependent DSB frequencies at replication fork barriers and revealed several orders of magnitude differences in DSB frequencies. Such high variability in genome breakage highlights the importance of quantification and shows how challenging data interpretation would be without the normalization provided by qDSB-Seq.

## Acknowledgements

This research was supported by the NIH grant R01GM112131 to M.R. (Y.Z., N.D., B.F., J.N., R.Y. and M.R.), Polish National Science Centre grant to M.S. (2015/17/D/NZ2/03711), and Foundation for Polish Science grant TEAM/2016-2 to K.G. This work was also supported by Ligue contre le Cancer (Equipe labelisee), Agence Nationale pour la Recherche (ANR) and Institut National du Cancer (INCa) grants to P.P. (B.P., R.F. and P.P.), National Science Center grant 2016/21/B/ST6/01471 to N.D., and a training fellowship from the Gulf Coast Consortia on the Computational Cancer Biology Training Program (CPRIT Grant No. RP170593) to Y.Z. The authors are grateful to Heather Lander of the Sealy Center for Structural Biology and Molecular Biophysics at UTMB, for editorial services for the manuscript.

## Author contributions

M.R. conceived qDSB-Seq and supervised and coordinated the project. M.R., Y.Z. and A.B. wrote the manuscript, K.G. P.P., B.P and M.S. edited the manuscript. Y.Z. performed data analysis and developed software, N.D. performed initial data analysis and developed software. A.B., K.G., B.P., P.P. and M.R. designed experiments. A.B. and M.S. performed i-BLESS and qDSB-Seq experiments. B.P., and R.F. prepared cells. Y.Z. prepared figures. R.Y., B.F. and J.N. contributed to software development and data analysis. M.S. performed library preparation and next-generation sequencing. All authors read the manuscript.

## ONLINE METHODS

### Strains and growth conditions

Yeast strains used in this study are listed in **Supplementary Table 5**. Cells were grown in YPD medium at 25°C until early log phase and were then arrested in G_1_ for 170 min with 8 µg/ml α-factor. For exposure to Zeocin cells were treated with 100 µg/ml Zeocin (Invivogen) for 1 hour. The I-SceI strain was cultured in YPR medium, galactose was added for 2 h to induce I-SceI cutting. For exposure to hydroxyurea, cells were released from G_1_ arrest by addition of 75 µg/ml Pronase (Sigma) and 200 mM HU was added 20 min before Pronase release followed by 1 h incubation. Collected cells were washed with cold SE buffer (5M NaCl, 500 mM EDTA, pH 7.5) and immediately subjected to DSB labeling.

### DSB sequencing

DSB labeling was performed using our i-BLESS method as described in^15^. Zeocin treated cells were additionally subjected to reaction with NEBNext® FFPE DNA Repair Mix prior to proximal adapter ligation. Sequencing libraries for i-BLESS and respective gDNA samples were prepared using ThruPLEX DNA-seq Kit (Rubicon Genomics). i-BLESS libraries were prepared without prior fragmentation and further size selection. Quality and quantity of the libraries were assessed on a 2100 Bioanalyzer using HS DNA Kit, and on a Qubit 2.0 Fluorometer using Qubit dsDNA HS Assay Kit (Life Technologies). The libraries were sequenced (2×70bp) on Illumina HiSeq2500/HiSeq4000 platforms, according to our modified experimental and software protocols for generation of high-quality data from low-diversity samples^16^.

### qDSB-Seq with NotI, SrfI, AsiSI, and BamHI digestion

In addition to DSB sequencing, as described above, a digestion with a restriction enzyme was performed before DSB labeling. Samples were treated with NotI (NEB, Thermo Scientific), SrfI (NEB), AsiSI (NEB), or BamHI (Thermo Scientific) for 1 h at 37°C. The dose and incubation time of these restriction enzymes were listed in **Supplementary Table 6**.

### qDSB-Seq with I-SceI spike-in

For I-SceI spike-in we used a yeast strain (I-SceI strain) with GAL inducible I-SceI endonuclease and a single I-SceI cutting site integrated at the ADH4 locus on chromosome VII. To measure the cleavage efficiency of I-SceI, cell aliquots were taken pre-(RAFF) and 2 h post-(GAL) cleavage induction, and total genomic DNA was extracted. DNA was serially diluted and amplified for 25 cycles with primers spanning the I-SceI cutting site. Cleavage efficiency was inferred by comparing the amount of amplified DNA in GAL (cut) vs. RAFF (uncut) conditions. We used CASY Cell Counter (Roche Applied Science) to mix this spike-in with our sample of interest (wild-type cells with replication stress induced by hydroxyurea treatment) in proportion 2:98. The cutting ratio of the I-SceI endonuclease expressed in the I-SceI strain was estimated using an unmixed I-SceI strain and **Equation (1)** below.

### Quantitative PCR

To validate cutting efficiency for NotI, input gDNA was analyzed by real-time PCR using primers flanking a selected NotI site at chrI: 114016-114023 (forward: AGAGTTGGGAATGTGTGCCC, reverse: GGGCAGCAACACAAAGTGTC) and KAPA SYBR® FAST kit (Life Technologies). Four technical replicates using two different concentrations of input DNA were performed. We compared the amount of PCR product amplified in untreated (C) vs. NotI treated cells (T) by data analysis based on the ΔC_T_ method^35^, where the ΔC_T_ value was obtained by subtraction of the C_T_ value in sample C from the C_T_ value in sample T. Final cutting efficiency was calculated as mean efficiency for all dilutions according to the formula below:

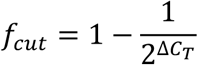

We used calibration data to empirically correct ΔC_T_

### Sequencing data analysis

We used *iSeq* (http://breakome.eu/software.html) to ensure sequencing data quality before mapping. Next, *iSeq* was used to remove i-BLESS proximal and distal barcodes (TCGAGGTAGTA and TCGAGACGACG, respectively). Reads labeled with the proximal barcode, which are directly adjacent to DSBs, were selected and mapped to the version of the yeast S288C genome sacCer3 (we manually corrected common polymorphisms) using bowtie^36^ v0.12.2 with the alignment parameters ‘-m1 –v1’ (to exclude ambiguous mapping and low-quality reads). For ribosomal DNA mapping in replication fork barrier analysis, we mapped sequencing reads using the parameter ‘-v1’ to allow multiple mapped reads. The end base pairs of the reads were trimmed using bowtie ‘-3’ parameter. The parameter choice was based on the *iSeq* quality report. For calculation of the absolute number of DSBs per cell only mapped reads were retained. Further, the reads identified as originating from telomere ends were removed. The telomeric reads were identified as those exhibiting the CAC motif in the whole AC-rich strand; regular expression C{0,3}AC{1,10} in the PERL language was used to identify them.

### Calculation of DSB frequencies per cell

Paired-end sequencing of gDNA or qPCR was used to measure the cutting efficiency of the endonuclease. For an enzyme with a single cutting site (e.g. I-SceI), we used the following procedure to calculate cutting efficiency (*f*_*cut*_) from whole genome paired-end sequencing data:

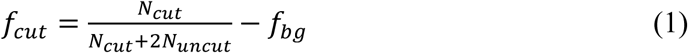

where, *N*_*cut*_ is the number of fragments cut by an enzyme, *N*_*uncut*_ is the number of uncut fragments covering the cutting site, and *f*_*bg*_ is the background level of breaks (e.g. resulting from sonication). *N*_*cut*_ fragments were counted in empirically determined, several nucleotide vicinities of the canonical cutting sites, based on visual examination of the read distribution. For enzyme with multiple cutting sites, reads mapped to each cutting site were first classified as “cut” or “uncut” and the results were summed over all cutting sites:

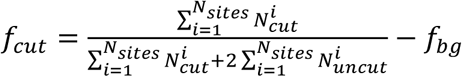

To estimate cutting efficiency, we used only cutting sites to which > 100 paired-end reads were mapped and their cutting efficiency was larger than 0. To estimate *f*_*bg*_, we randomly selected genomic windows of the same size as those used to count cut and uncut fragments and estimated “cutting efficiency” in those intervals using the left part of **Equation (1)**. For clarity, these errors are omitted in **Equations (2)** to (**4)**.

Next, we calculated the number of spike-in DSBs induced at restriction sites, *B*_*cut*_:

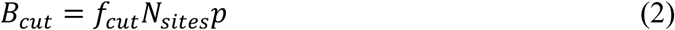

where *f*_*cut*_ is the cutting efficiency in undiluted samples, *N*_*sites*_ is the number of used enzyme restriction sites (e.g. 39 for NotI) and *p* is the proportion of digested cells (*p* = 1 unless mixing with an *in vivo* digested construct is used).

Then we computed the number of mapped sequencing reads per DSB or the coefficient, *α*:

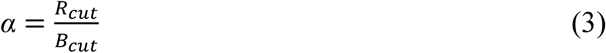

where *R*_*cut*_ is the number of labeled reads mapped to the cutting sites and *B*_*cut*_ is the total number of induced DSBs.

Finally, we computed studied DSBs per cell (*B*_*studied*_) using the following formula:

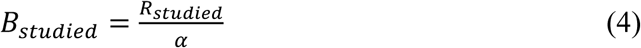

where *B*_*studied*_ is the number of studied DSBs per cell in the whole genome, or in a specific region (eg. a replication region), or at a specific location (eg. an enzyme cutting site). In this study, we calculated the studied breaks per cell for the whole genome after subtracting reads generated from enzyme cutting sites, telomeres, and ribosomal DNA. Errors for *B*_*studied*_ are the standard deviation of breaks calculated from different cutting sites for enzymes with multiple cutting sites (**Supplementary Table 1**). Based on replicates, we concluded that thus calculated errors are conservatively estimated. For an enzyme with a single cutting site in a given genome, errors for *B*_*cell*_ were assigned using computed errors of the cutting efficiencies from *f*_*bg*_.

### Background estimation and removal

To quantify DSBs likely resulting from broken forks near origins, we first removed background not related to replication. To define such background, we calculated DSB density in a 500 bp sliding window with a 50 bp step; the peak of this distribution was assumed to be background DSB frequency. This background was subtracted from the data at each position, resulting negative values were assigned to zero.

### Analysis of fragile regions and enrichment

Hygestat_BLESS v1.2.3 in the *iSeq* package (http://breakome.eu/software.html) was used to identify fragile regions (i.e. regions with significant increase of the read numbers in treatment versus control samples), which were defined using the hypergeometric probability distribution and Benjamini-Hochberg correction. To evaluate the enrichment of fragile regions on nucleosomes, we used hygestat_annotations v2.0, which computed the proportion of mappable nucleotides belonging to both the fragile regions and the nucleosomes, and the proportion of mappable nucleotides belonging to both genomic regions and the nucleosomes. To estimate the p-value for the feature enrichment inside fragile regions, we used 1000 permutations to calculate the empirical distribution of the ratio under the null hypothesis.

### Estimation of one-ended DSBs

To estimate the total number of one-ended DSBs, we performed hypergeometric test based on the number of i-BLESS sequencing reads from Watson and Crick strands using Hygestat_BLESS v1.2.3 in the *iSeq* package with a 500 nt window size. Regions with *P* < 1e-10 were classified as one-ended DSB regions, *P* value was corrected by the Bonferroni correction. The subtraction between reads from Watson and Crick was treated as the number of one-ended reads used to calculate one-ended DSBs using the DSB calculation method.

### Comparison of DSB levels between ZEO and G_1_ samples

We used read counts for 5000 nt mappable intervals produced by hygestat_BLESS; ZEO read numbers were normalized using qDSB-Seq quantification. We evaluated the null hypothesis that the number of DSBs in G_1_ cells is the same or lower than in ZEO using very conservative 5 standard deviation confidence intervals (assuming Poisson distribution of reads). All genomic windows with >17 reads in 5 kb were significantly enriched in DSBs in ZEO as compared with G_1_ cells (*P* < 2e-12, calculated using the hypergeometric probability distribution and the Bonferroni correction).

### DNA secondary structure and G-quadruplex prediction

DNA secondary structures were defined by free energy at 37°C using UNAFold^37^ v3.8 in a 50 bp sliding window with a 25 bp step along the whole yeast genome. We predicted G-quadruplexes (both canonical intrastrand and non-canonical inter-strand) in the budding yeast genome using AllQuads^38^ software, with the standard 7-nt threshold on loop length.

### Statistical analysis

Results of quantification are shown as mean ± s.d. To conduct enrichment analysis, the *P* values were first calculated using the hypergeometric distribution function as implemented in the GNU Scientific Library for C++ and then corrected for multiple hypothesis testing using the Benjamini-Hochberg method. The threshold for statistical significance was *P* < 0.05.

### Code availability

Custom code used in this study is available upon request from authors or http://breakome.eu/software.html.

### Data availability

The DSB sequencing data will be available upon publication at Sequence Read Archive.

## References

1. Khanna, K.K. & Jackson, S.P. DNA double-strand breaks: signaling, repair and the cancer connection. Nature genetics 27, 247–254 (2001).

2. Slaymaker, I.M. et al. Rationally engineered Cas9 nucleases with improved specificity. Science 351, 84–88 (2016).

3. Crosetto, N. et al. Nucleotide-resolution DNA double-strand break mapping by next-generation sequencing. Nat Methods 10, 361–365 (2013).

4. Hoffman, E.A., McCulley, A., Haarer, B., Arnak, R. & Feng, W. Break-seq reveals hydroxyurea-induced chromosome fragility as a result of unscheduled conflict between DNA replication and transcription. Genome Res 25, 402–412 (2015).

5. Lensing, S.V. et al. DSBCapture: in situ capture and sequencing of DNA breaks. Nature methods 13, 855–857 (2016).

6. Canela, A. et al. DNA Breaks and End Resection Measured Genome-wide by End Sequencing. Molecular cell 63, 898–911 (2016).

7. Yan, W.X. et al. BLISS is a versatile and quantitative method for genome-wide profiling of DNA double-strand breaks. Nature communications 8, 15058 (2017).

8. Ran, F.A. et al. In vivo genome editing using Staphylococcus aureus Cas9. Nature 520, 186–191 (2015).

9. Aymard, F. et al. Genome-wide mapping of long-range contacts unveils clustering of DNA double-strand breaks at damaged active genes. Nature structural & molecular biology 24, 353–361 (2017).

10. Popp, H.D., Brendel, S., Hofmann, W.K. & Fabarius, A. Immunofluorescence Microscopy of gammaH2AX and 53BP1 for Analyzing the Formation and Repair of DNA Double-strand Breaks. J Vis Exp (2017).

11. Chailleux, C. et al. Quantifying DNA double-strand breaks induced by site-specific endonucleases in living cells by ligation-mediated purification. Nat Protoc 9, 517–528 (2014).

12. Gregoire, M.C. et al. Quantification and genome-wide mapping of DNA double-strand breaks. DNA repair 48, 63–68 (2016).

13. Clement, K., Farouni, R., Bauer, D.E. & Pinello, L. AmpUMI: design and analysis of unique molecular identifiers for deep amplicon sequencing. Bioinformatics 34, i202–i210 (2018).

14. Smith, T., Heger, A. & Sudbery, I. UMI-tools: modeling sequencing errors in Unique Molecular Identifiers to improve quantification accuracy. Genome Res 27, 491–499 (2017).

15. Biernacka, A. et al. i-BLESS is an ultra-sensitive method for detection of DNA double-strand breaks. Commun Biol 1, 181 (2018).

16. Mitra, A., Skrzypczak, M., Ginalski, K. & Rowicka, M. Strategies for achieving high sequencing accuracy for low diversity samples and avoiding sample bleeding using illumina platform. PLoS One 10, e0120520 (2015).

17. Lobrich, M. et al. gammaH2AX foci analysis for monitoring DNA doublestrand break repair: strengths, limitations and optimization. Cell Cycle 9, 662–669 (2010).

18. Thongsroy, J. et al. Replication-independent endogenous DNA double-strand breaks in Saccharomyces cerevisiae model. PLoS One 8, e72706 (2013).

19. Shimada, K. et al. TORC2 signaling pathway guarantees genome stability in the face of DNA strand breaks. Molecular cell 51, 829–839 (2013).

20. Lee, W. et al. A high-resolution atlas of nucleosome occupancy in yeast. Nature genetics 39, 1235–1244 (2007).

21. Povirk, L.F. DNA damage and mutagenesis by radiomimetic DNA-cleaving agents: bleomycin, neocarzinostatin and other enediynes. Mutat Res 355, 71–89 (1996).

22. Bochman, M.L., Paeschke, K. & Zakian, V.A. DNA secondary structures: stability and function of G-quadruplex structures. Nature reviews. Genetics 13, 770–780 (2012).

23. Hershman, S.G. et al. Genomic distribution and functional analyses of potential G-quadruplex-forming sequences in Saccharomyces cerevisiae. Nucleic acids research 36, 144–156 (2008).

24. Koc, A., Wheeler, L.J., Mathews, C.K. & Merrill, G.F. Hydroxyurea arrests DNA replication by a mechanism that preserves basal dNTP pools. The Journal of biological chemistry 279, 223–230 (2004).

25. Singh, A. & Xu, Y.J. The Cell Killing Mechanisms of Hydroxyurea. Genes (Basel) 7 (2016).

26. Sasaki, M. & Kobayashi, T. Ctf4 Prevents Genome Rearrangements by Suppressing DNA Double-Strand Break Formation and Its End Resection at Arrested Replication Forks. Molecular cell 66, 533–545 e535 (2017).

27. Kobayashi, T. The replication fork barrier site forms a unique structure with Fob1p and inhibits the replication fork. Molecular and cellular biology 23, 9178–9188 (2003).

28. Kobayashi, T., Horiuchi, T., Tongaonkar, P., Vu, L. & Nomura, M. SIR2 regulates recombination between different rDNA repeats, but not recombination within individual rRNA genes in yeast. Cell 117, 441–453 (2004).

29. Weitao, T., Budd, M. & Campbell, J.L. Evidence that yeast SGS1, DNA2, SRS2, and FOB1 interact to maintain rDNA stability. Mutat Res 532, 157–172 (2003).

30. Burkhalter, M.D. & Sogo, J.M. rDNA enhancer affects replication initiation and mitotic recombination: Fob1 mediates nucleolytic processing independently of replication. Molecular cell 15, 409–421 (2004).

31. Weitao, T., Budd, M., Hoopes, L.L. & Campbell, J.L. Dna2 helicase/nuclease causes replicative fork stalling and double-strand breaks in the ribosomal DNA of Saccharomyces cerevisiae. The Journal of biological chemistry 278, 22513–22522 (2003).

32. Di Felice, F., Cioci, F. & Camilloni, G. FOB1 affects DNA topoisomerase I in vivo cleavages in the enhancer region of the Saccharomyces cerevisiae ribosomal DNA locus. Nucleic acids research 33, 6327–6337 (2005).

33. Zhu, Y., Biernacka, A., Pardo, B., Forey, R., Dojer, N., Yousefi, R., Nde, J., Fongang, B., Mitra, A., Li, J., Skrzypczak, M., Kudlicki, A., Pasero, P., Ginalski, K., Rowicka, M. Integrated analysis of patterns of DNA breaks reveals break formation mechanisms and their population distribution during replication stress. BioRxiv (2017).

34. Caron, P. et al. Non-redundant Functions of ATM and DNA-PKcs in Response to DNA Double-Strand Breaks. Cell Rep 13, 1598–1609 (2015).

35. Schmittgen, T.D. & Livak, K.J. Analyzing real-time PCR data by the comparative C-T method. Nat Protoc 3, 1101–1108 (2008).

36. Langmead, B., Trapnell, C., Pop, M. & Salzberg, S.L. Ultrafast and memory-efficient alignment of short DNA sequences to the human genome. Genome Biol 10, R25 (2009).

37. Markham, N.R. & Zuker, M. DINAMelt web server for nucleic acid melting prediction. Nucleic acids research 33, W577–581 (2005).

38. Kudlicki, A.S. G-Quadruplexes Involving Both Strands of Genomic DNA Are Highly Abundant and Colocalize with Functional Sites in the Human Genome. PLoS One 11, e0146174 (2016).

39. Yabuki, N., Terashima, H. & Kitada, K. Mapping of early firing origins on a replication profile of budding yeast. Genes to cells: devoted to molecular & cellular mechanisms 7, 781–789 (2002).

