## Supplementary Materials for "qDSB-Seq: quantitative DNA double-strand break sequencing"

### Supplementary Figures

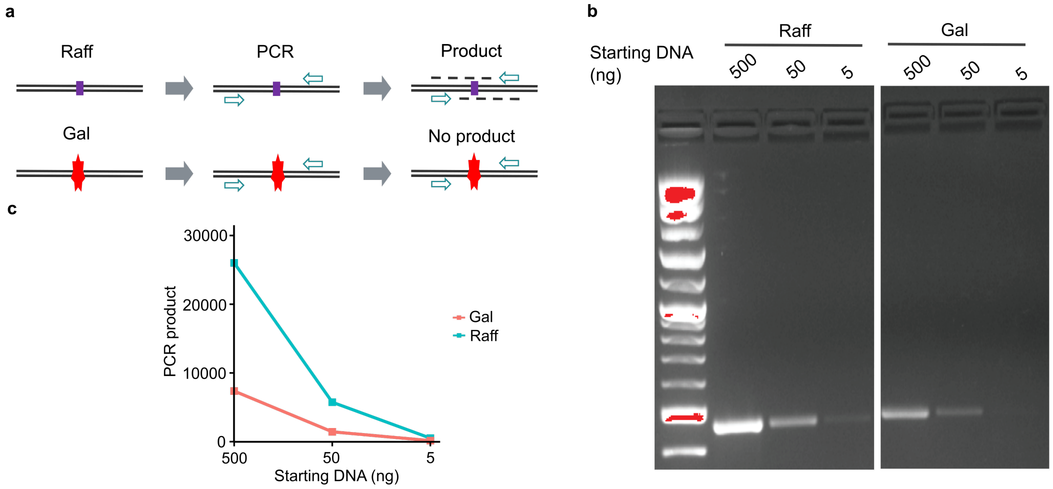


**Supplementary Figure 1**. Validation of cutting efficiency using semi-quantitative PCR. **(a)** A schema for sqPCR design and its results for cells cultured in raffinose (no I-SceI digestion) and galactose (I-SceI digestion). The purple rectangle is I-SceI recognition site. The star symbol represents the digestion of I-SceI recognition sequence induced by galactose. **(b)** Gel electrophoresis of PCR products in Raff and Gal samples for different amount of starting DNA. **(c)** Quantification of PCR products using agarose gel.

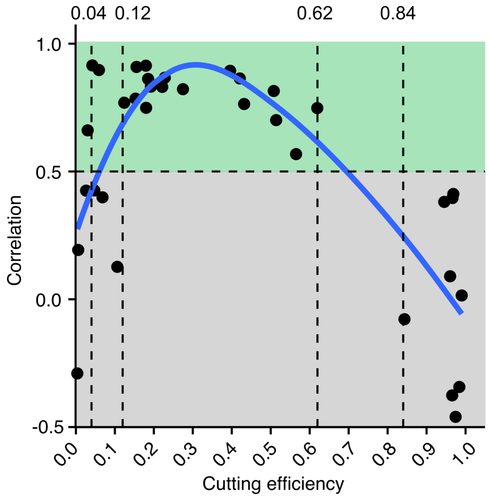


**Supplementary Figure 2.** Pearson correlation between cutting efficiencies and the labeled reads for all sites recognized by a given enzyme was calculated for 25 samples treated with restriction enzymes with multiple cutting sites (35 separate digestions). The green and grey areas were divided by correlation 0.5 that shows strong relationship between cutting efficiencies and the labeled reads for quantification.

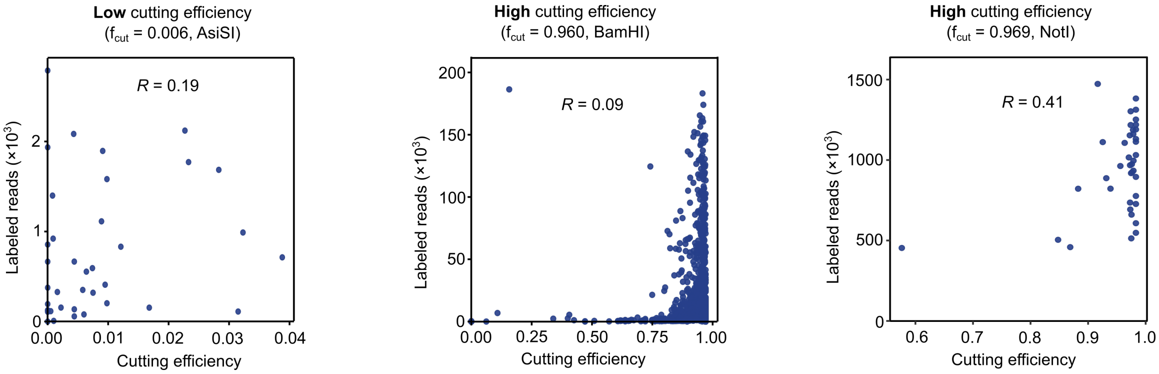


**Supplementary Figure 3.** Employment of extremely low or high cutting efficiencies results in loss of proportional relationship between the labeled reads and cutting efficiencies at enzyme cutting sites. Pearson correlation ($R$) between cutting efficiencies and the labeled reads was calculated for all sites recognized by a given enzyme. From left to right, AsiSI, BamHI, NotI treated samples are shown.

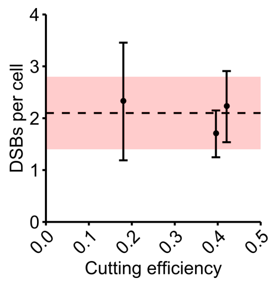


**Supplementary Figure 4.** Quantification of DSBs in $G_1$ *pif1-m2* mutant cells. The dashed lines and the stripes are respectively the mean value and 95% confidence interval for all three samples; mean values with error bars for individual samples are also shown.

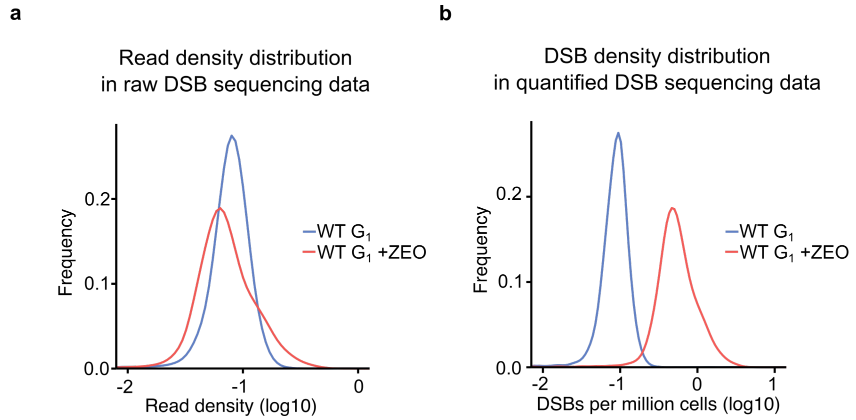


**Supplementary Figure 5.** Density plot of raw and quantified DSB sequencing data for Zeocin-treated and untreated $G_1$ samples. **(a)** The distribution of read density. Read density was defined as the number of i-BLESS reads mapped to a given region, divided by region length, here a 500 bp sliding window and a 50 bp step. **(b)** The distribution of DSB density. DSB density was calculated from DSBs per million cells using the same method in **(a)**.

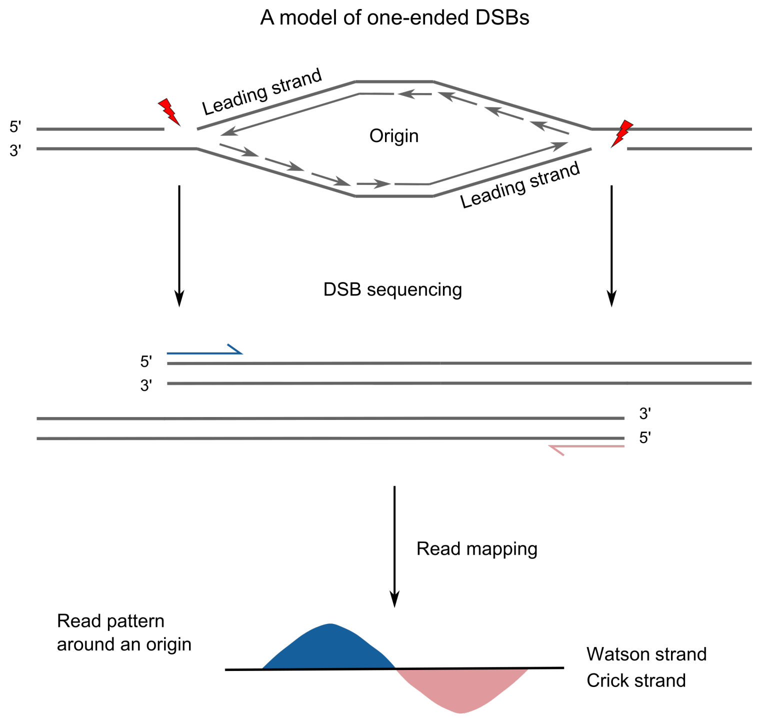


**Supplementary Figure 6.** A characteristic one-ended DSB pattern of i-BLESS reads. The pattern arises around replication origins due to broken replication forks resulting in one-ended DSBs.

### Supplementary Tables

**Supplementary Table 1.** Absolute DSB frequencies per cell and its variation influenced by low and high cutting efficiencies in $G_1$ untreated samples. To measure the variation, standard deviation (s.d.) of DSBs was calculated by individual cutting sites of NotI, SrfI, AsiSI, and BamHI. Only effective cutting sites were used for calculation as described in Methods.

| Sample name | Enzyme | Average Cutting efficiency | Studied DSBs | S.d. of studied DSBs | S.d. of studied DSBs / Studies DSBs | Number of enzyme cutting sites per Mbases | % of enzyme-induced spike-in reads |
|---|---|---|---|---|---|---|---|
| **Adequate cutting efficiency** | | | | | | | |
| $G_1$ exp6 | NotI | 0.04 | 0.7 | 0.6 | 0.84 | 3 | 20 |
| $G_1$ exp6 | SrfI | 0.06 | 0.8 | 0.4 | 0.49 | 2 | 11 |
| $G_1$ exp2 | SrfI | 0.07 | 0.6 | 0.2 | 0.41 | 2 | 20 |
| $G_1$ exp3 | SrfI | 0.11 | 1.7 | 0.7 | 0.44 | 2 | 20 |
| $G_1$ exp1 | NotI | 0.18 | 1.1 | 0.3 | 0.25 | 3 | 87 |
| Mean | | | 1.0 | | | | |
| S.d. | | | 0.4 | | | | |
| $G_1$ pif1 exp1 | NotI | 0.18 | 2.3 | 1.1 | 0.49 | 3 | 75 |
| $G_1$ pif1 exp2 | NotI | 0.40 | 1.7 | 0.5 | 0.27 | 3 | 90 |
| $G_1$ pif1 exp3 | NotI | 0.42 | 2.2 | 0.7 | 0.31 | 3 | 88 |
| Mean | | | 2.1 | | | | |
| S.d. | | | 0.3 | | | | |
| **Low cutting efficiency** | | | | | | | |
| $G_1$ exp3 | AsiSI | 0.004 | 6774 | 1770 | 0.26 | 3 | 0 |
| $G_1$ exp2 | AsiSI | 0.006 | 39 | 375 | 9.65 | 3 | 0 |
| $G_1$ exp4 | AsiSI | 0.03 | 9.5 | 16.0 | 1.69 | 3 | 1 |
| Mean | | | 2274 | | | | |
| S.d. | | | 3897 | | | | |
| **High cutting efficiency** | | | | | | | |
| $G_1$ exp4 | SrfI | 0.84 | 1.3 | 0.5 | 0.38 | 2 | 57 |
| $G_1$ exp5 | NotI | 0.95 | 4.0 | 1.9 | 0.47 | 3 | 22 |
| $G_1$ exp15 | BamHI | 0.96 | 2.2 | 27 | 12.40 | 139 | 93 |
| $G_1$ exp5 | AsiSI | 0.97 | 7.9 | 17 | 2.20 | 3 | 22 |
| $G_1$ exp3 | NotI | 0.97 | 11 | 4.1 | 0.38 | 3 | 62 |
| $G_1$ exp2 | NotI | 0.97 | 5.6 | 1.9 | 0.33 | 3 | 69 |
| $G_1$ exp6 | AsiSI | 0.98 | 5.2 | 2.5 | 0.48 | 3 | 57 |
| $G_1$ exp5 | SrfI | 0.98 | 4.0 | 0.9 | 0.22 | 2 | 13 |
| $G_1$ exp4 | NotI | 0.99 | 6.9 | 3.4 | 0.49 | 3 | 31 |
| Mean | | | 5.3 | | | | |
| S.d. | | | 2.9 | | | | |

**Supplementary Table 2.** Pearson correlation between the i-BLESS labeled reads and cutting efficiencies for all sites recognized by a given enzyme. The average cutting efficiency was calculated from all cutting sites digested by an enzyme; the results are sorted according to descending R.

| **Enzyme** | **Sample name** | **Average cutting efficiency** | ***R*** |
|---|---|---|---|
| NotI | $G_1$ exp6 | 0.04 | 0.915 |
| NotI | $G_1$ exp1 | 0.18 | 0.914 |
| NotI | ZEO | 0.16 | 0.909 |
| SrfI | $G_1$ exp6 | 0.06 | 0.897 |
| NotI | $G_1$ pif1 exp2 | 0.40 | 0.894 |
| NotI | S exp4 | 0.23 | 0.867 |
| NotI | $G_1$ pif1 exp3 | 0.42 | 0.864 |
| NotI | S exp1 | 0.18 | 0.862 |
| NotI | S exp2 | 0.19 | 0.832 |
| NotI | $G_1$ exp7 | 0.22 | 0.831 |
| NotI | S exp6 | 0.28 | 0.822 |
| NotI | $G_1$ exp10 | 0.51 | 0.815 |
| NotI | WT S | 0.15 | 0.786 |
| NotI | WT +CPT | 0.12 | 0.769 |
| NotI | $G_1$ exp8 | 0.43 | 0.764 |
| NotI | $G_1$ pif1 exp1 | 0.18 | 0.749 |
| NotI | WT +HU | 0.62 | 0.748 |
| NotI | $G_1$ exp9 | 0.51 | 0.701 |
| AsiSI | $G_1$ exp4 | 0.03 | 0.661 |
| NotI | $G_1$ exp7 | 0.56 | 0.568 |
| NotI | S exp5 | 0.05 | 0.426 |
| NotI | S exp3 | 0.03 | 0.425 |
| NotI | $G_1$ exp2 | 0.97 | 0.412 |
| SrfI | $G_1$ exp2 | 0.07 | 0.399 |
| NotI | $G_1$ exp3 | 0.97 | 0.396 |
| NotI | $G_1$ exp5 | 0.94 | 0.381 |
| AsiSI | $G_1$ exp2 | 0.006 | 0.193 |
| SrfI | $G_1$ exp3 | 0.11 | 0.127 |
| BamHI | BamHI | 0.96 | 0.090 |

| | | | |
|---|---|---|---|
| NotI | $G_1$ exp4 | 0.99 | 0.015 |
| SrfI | $G_1$ exp4 | 0.84 | -0.078 |
| AsiSI | $G_1$ exp3 | 0.004 | -0.290 |
| SrfI | $G_1$ exp5 | 0.98 | -0.343 |
| AsiSI | $G_1$ exp5 | 0.97 | -0.376 |
| AsiSI | $G_1$ exp6 | 0.98 | -0.460 |

**Supplementary Table 3.** DSB frequencies per cell near Replication Fork Barriers (RFBs) in a rDNA array.

| | rDSB-1 | rDSB-2 | rDSB-3 | DSBs on RFBs | Std. DSBs on RFBs |
|---|---|---|---|---|---|
| WT $G_1$ | 0.00 | 0.00 | 0.00 | 0.00 | 0.00 |
| WT S | 0.80 | 0.30 | 0.00 | 1.09 | 0.23 |
| WT +CPT | 0.76 | 0.28 | 0.11 | 1.15 | 0.28 |

**Supplementary Table 4.** Restriction enzymes cutting sites for *Saccharomyces cerevisiae*, *Homo Sapiens*, *Mus musculus*, *Drosophila melanogaster*, *Arabidopsis thaliana*, and *Caenorhabditis elegans* (separate file due to size).

**Supplementary Table 5.** Yeast strains used in this study.

| Strain | Genotype |
|---|---|
| YBP-275 (I-SceI) | *MATa-inc, bar1Δ, ade2-1, can1-100, leu2::SFA1, trp1-1, ura3-1, lys2::GAL1p-ISCEI, adh4::URA3::GAL1p::leu2Δ3'::ACT1iΔ3'::IsceIsite his3::HYG:HOsite::ACT1-iΔ5'::leu2Δ5'* |
| WT | *MATa, ade2-1, trp1-1, can1-100, leu2-3,112, his3-11,15, ura3; GAL, psi+, RAD5; URA3::GPD-TK(7x)* |
| *pif1-m2* | *MATa, ade2-1, ura3-1, his3-11,15, leu2-3, 112, trp1-1, CAN1, GAL, PSI+, sml1:TRP1, pif1-m2* |

**Supplementary Table 6.** Dose and incubation time of restriction enzymes.

| Sample name | Cell cycle phase | Enzyme | Enzyme dose | Incubation time |
|---|---|---|---|---|
| $G_1$ exp1 | $G_1$ | NotI | 12.5 nl/ml* | 60 min |
| $G_1$ exp2 | $G_1$ | NotI | 250 U | 60 min |
| $G_1$ exp2 | $G_1$ | AsiSI | 250 U | 60 min |
| $G_1$ exp2 | $G_1$ | SrfI | 250 U | 60 min |
| $G_1$ exp3 | $G_1$ | NotI | 250 U | 60 min |
| $G_1$ exp3 | $G_1$ | AsiSI | 250 U | 60 min |
| $G_1$ exp3 | $G_1$ | SrfI | 250 U | 60 min |
| $G_1$ exp4 | $G_1$ | NotI | 250 U | 60 min |
| $G_1$ exp4 | $G_1$ | AsiSI | 250 U | 60 min |
| $G_1$ exp4 | $G_1$ | SrfI | 250 U | 60 min |
| $G_1$ exp5 | $G_1$ | NotI | 250 U | 60 min |
| $G_1$ exp5 | $G_1$ | AsiSI | 250 U | 60 min |
| $G_1$ exp5 | $G_1$ | SrfI | 250 U | 60 min |
| $G_1$ exp6 | $G_1$ | NotI | 1 U | 60 min |
| $G_1$ exp6 | $G_1$ | AsiSI | 250 U | 120 min |
| $G_1$ exp6 | $G_1$ | SrfI | 1 U | 60 min |
| $G_1$ exp7 | $G_1$ | NotI | 5 nl/ml | 60 min |
| $G_1$ exp8 | $G_1$ | NotI | 5 nl/ml | 60 min |
| $G_1$ exp9 | $G_1$ | NotI | 5 nl/ml | 60 min |
| $G_1$ exp10 | $G_1$ | NotI | 5 nl/ml | 60 min |
| $G_1$ exp11 | $G_1$ | NotI | 5 nl/ml | 60 min |
| $G_1$ pif1 exp1 | $G_1$ | NotI | 5 nl/ml | 60 min |
| $G_1$ pif1 exp3 | $G_1$ | NotI | 5 nl/ml | 60 min |
| $G_1$ pif1 exp2 | $G_1$ | NotI | 5 nl/ml | 60 min |
| $G_1$ exp15 | $G_1$ | BamHI | 5 µl/ml | 60 min |
| ZEO | $G_1$ | NotI | 12.5 nl/ml | 60 min |
| WT S | S | NotI | 5 nl/ml | 60 min |
| WT +CPT | S | NotI | 5 nl/ml | 60 min |
| WT +HU | S | NotI | 5 nl/ml | 60 min |
| S exp1 | S | NotI | 5 nl/ml | 60 min |
| S exp2 | S | NotI | 5 nl/ml | 60 min |
| S exp3 | S | NotI | 5 nl/ml | 60 min |
| S exp4 | S | NotI | 5 nl/ml | 60 min |
| S exp5 | S | NotI | 5 nl/ml | 60 min |
| S exp6 | S | NotI | 5 nl/ml | 60 min |

Note: * nl/ml: nanoliter of enzyme solution per milliliter of reaction.
